# The mechano-chemistry of a monomeric reverse transcriptase

**DOI:** 10.1101/194159

**Authors:** Omri Malik, Hadeel Khamis, Sergei Rudnizky, Ariel Kaplan

## Abstract

Retroviral reverse transcriptase catalyses the synthesis of an integration-competent dsDNA molecule, using as a substrate the viral RNA. Using optical tweezers, we follow the Murine Leukemia Virus reverse transcriptase as it performs strand-displacement polymerization on a template under mechanical force. Our results indicate that reverse transcriptase functions as a Brownian ratchet, with dNTP binding as the rectifying reaction of the ratchet. We also found that reverse transcriptase is a relatively passive enzyme, able to polymerize on structured templates by exploiting their thermal breathing. Finally, our results indicate that the enzyme enters the recently characterized backtracking state from the pre-translocation complex.

## INTRODUCTION

The virally-encoded reverse transcriptase (RT) is responsible for genome replication in retroviruses. RT is a DNA-and RNA-dependent DNA polymerase that catalyses the synthesis of an integration-competent dsDNA molecule. RTs have the “right hand” structure typical of cellular DNA polymerases, with palm, fingers and thumb subdomains, and also a spatially separated RNase H domain(1). Using both steady state and pre-steady state techniques(2–7), previous studies have described a mechanism of polymerization by RT, which is similar to the mechanism of other DNA polymerases. However, the rates of polymerization by RTs are fairly slow and their processivity is poor relative to other DNAPs (8). In addition, as opposed to most (but not all(9–13)) DNAPs, which require a single strand DNA (ssDNA) template, RT is capable of efficiently unwinding duplexes in the template during polymerization. This strand displacement (SD) synthesis activity by RT is required for the polymerization on the highly structured vRNA(14), the removal of RNase H resistant RNA fragments(15) and polymerization on the long terminal repeats(16). Importantly, although the rates of both displacement and non-displacement DNA synthesis vary for different sites over the template, SD synthesis is, on the average, slower by a factor of 3–4(17), as compared to primer extension synthesis.

Remarkably, although numerous structural and biochemical analyses have been performed to elucidate the mechanisms governing dNTP binding(18, 19), induced conformational change(20–22) and catalysis(23, 24), much less is known about the mechanisms and kinetics of RT’s translocation. The first question that needs to be answered in order to incorporate the translocation step into the enzyme’s cycle, pertains to the location of the translocation step in the order of events that comprise a nucleotide addition cycle. Single molecule methods have been implemented to dissect the mechano-chemical cycle of RNA and DNA polymerases by using mechanical force to assist or inhibit translocation(25). For instance, the location of the translocation step for Phi29 DNA polymerase and Escherichia coli RNAP was found to take place after pyrophosphate (PPi) release and prior to dNTP binding(13, 26). Next, the *mechanism* of translocation needs to be elucidated. Two alternative mechanisms have been proposed: a “power stroke” (PS) mechanism, where a certain chemical step (dNTP hydrolysis or PPi release) directly powers translocation, and a “Brownian Ratchet” (BR) mechanism, where the enzyme moves back and forth between the pre- and post-translocation state, driven by thermal energy, and a chemical reaction traps the post-translocation configuration, driving the reaction forward. Interestingly, despite the paucity of experimental data, some controversy exists regarding the mechanism of translocation by RT. An active PS model for HIV-1 RT is supported by previous studies showing that nucleotide binding is associated with displacement of the highly conserved YMDD motif that is part of the enzymes active site(27). Release of this “loaded spring” following catalysis may provide the drive for the translocation reaction. In contrast, studies using a site-specific footprinting technique that allows distinguishing between the pre- and post-translocated states(28) support a BR mechanism: Marchand et. al. reported that the efficiency of inhibition of HIV-1 RT by the PPi analogue phosphonoformic acid (Foscarnet) correlates with the presence of sequences that favor the pre-translocation state, suggesting that this state is accessible during the incorporation cycle, and arguing against a PS model(29). Their results suggest that the pre- and post-translocation states equilibrate immediately following each nucleotide incorporation event, and that the presence of the templated nucleotide can stabilize and trap the post-translocated complex. This type of mechanism was demonstrated for RNA polymerase(30), as well as for the replicative Phi 29 DNA polymerase(13). Finally, given that RT is not stalled by duplexes in its template, it is necessary to clarify how RT copes with the obstacle presented by the duplex DNA. Generally, unwinding of nucleic-acid duplexes by enzymes, such as helicases, has been described by two extreme models(31): In an “active” model the enzyme unwinds the nucleic acid duplex by directly destabilizing the hairpin fork. In contrast, in the “passive” model, the enzyme is unable to destabilize the duplex ahead, but it exploits the spontaneous, thermally-driven opening of the duplex (thermal “breathing”). These are extreme mechanisms, and most enzymes will share some of the properties of both, i.e. will partially destabilize the duplex ahead, but will still be affected by the duplex spontaneous breathing(32, 33). Taken together, the location of the translocation step, the mechanism of translocation, and the mechanism of duplex unwinding determine how RT responds to changes in [dNTP] and to the presence and magnitude of an external force. Hence, experimentally characterizing these responses can shed light on the mechano-chemistry of RT.

Here, we use optical tweezers to elucidate the mechanism of translocation of the Moloney Murine Leukemia Virus (M-MLV) RT. By following strand-displacement polymerization by RT on a DNA hairpin, and separating active polymerization phases from pauses, we were able to characterize the force- and nucleotide-dependence of RT’s velocity, and compare these dependencies with the predictions of detailed kinetic models. Furthermore, we show that a comparison of the experimentally measured density of pauses with the predictions of the kinetic models provides a powerful tool to discriminate between plausible models, identify the kinetic step from which RT enters the pauses, and calculate the intrinsic rate of pausing by RT.

## MATERIALS AND METHODS

### Molecular constructs for single molecule experiments

The DNA sequences used as a polymerization template (Supplementary Table 1) was amplified by PCR from a plasmid that was a generous gift from Daniela Rhodes (MRC, Cambridge, UK)). Primers used for the amplification reactions are listed in Supplementary Table 2. The constructs were digested using DraIII-HF (New England Biolabs) overnight according to the manufacturer’s instructions. A 10 bp hairpin (Sigma) was ligated to the construct using T4 DNA ligase (New England Biolabs), in a reaction with 1:10 molar excess of the hairpin, at 16 °C. The construct was subsequently digested overnight with BglI (New England Biolabs). Two 600 bp DNA handles, each incorporating a specific tag (digoxigenin and biotin), were generated using commercially purchased 5’ modified primers (Supplementary Table 3) in a standard PCR reaction. Two of the primers were designed to contain repeats of three DNA sequences recognized by single strand nicking enzymes: Nt.BbvCI and Nb.BbvCI (both from New England Biolabs) on the biotin-tagged handle and on the digoxigenin-tagged handle, respectively. The nicking enzymes generated 29 nt complementary overhangs on each handle. Handles were mixed at equal molar ratios for DNA annealing, creating a ~1200 bp fragment of annealed DNA handles. The polymerization templates were ligated to the DNA handles using a rapid ligase system (Promega) in 3:1 molar ratio, 30 min at room temp.

### Reagents

M-MLV RT and dNTPs were purchased from New England Biolabs and Sigma, respectively.

### Optical Tweezers

Experiments were performed in a custom-made double-trap optical tweezers apparatus, as previously described(34, 35). Briefly, the beam from a 852 nm laser (TA PRO, Toptica) was coupled into a polarization-maintaining single-mode optical fiber. The collimated beam out of the fiber, with a waist of w_0_=4mm, was split by a polarizing beamsplitter into two orthogonal polarizations, each directed into a mirror and combined again with a second beamsplitter. One of the mirrors is mounted on a nanometer scale mirror mount (Nano-MTA, Mad City Labs). A X2 telescope expands the beam, and also images the plane of the mirrors into the back focal plane of the focusing microscope objective (Nikon, Plan Apo VC 60X, NA/1.2). Two optical traps are formed at the objective’s focal plane, each by a different polarization, and with a typical stiffness of 0.3-0.5 pN/nm. The light is collected by a second, identical objective, the two polarizations separated by a beamsplitter, and imaged onto two Position Sensitive Detectors (First Sensor). The position of the beads relative to the center of the trap is determined by back focal plane interferometry(36). Calibration of the setup was done by analysis of the thermal fluctuations of the trapped beads(37), which were sampled at 100kHz.

### Single molecule polymerization experiments

The full polymerization construct was incubated for 15 min on ice with 0.9 µm polystyrene beads (Spherotech), coated with streptavidin (SA), and diluted 1000-fold in RT buffer (RTB; 50 mM Tris-HCl, 75 mM KCl, 3 mM MgCl2, 10 mM DTT, pH 8.3 @ 25°C). The SA beads bound to the DNA constructs, together with 0.9 µm anti-digoxigenin (*α*D) coated beads were introduced into the microfluidic channel filled with RTB. Tether formation was performed *in situ* (inside the experimental chamber) by trapping a SA coated bead in one trap, trapping an *α*D bead in the second trap, and bringing the two beads into close proximity to allow binding of the digoxigenin tag in the DNA to the *α*D in the bead. After a few seconds, the beads were moved away from each other while monitoring changes in the force. Establishment of a tether is indicated by an increase in force as the traps are separated. The polymerization reaction was initiated by flowing activity buffer (RTB with the addition of 0.5-250 µM dNTP and 200U/ml RT) into the chamber.

Experiments were conducted in a semi-passive mode(38), in which polymerization takes place with no feedback on the force but where, if the force exceeds a predetermined value, the position of the steerable trap is rapidly changed in a single step and in a direction and magnitude that are expected to restore the measured force to the range of forces that were pre-established (typically, ±1.5 pN of the nominal force). As a result, our polymerization data consists of intervals of passive-mode operation, that are separated by sudden “jumps” in the position of the steerable trap. To detect these jumps, the voltage on the piezoelectric controlled mirror was sampled at 2.5 kHz and filtered with a 3^rd^ order Savitzky-Golay filter with a frame length of 101 points. Time points where the derivative of this filtered signal was larger than 0.5 V s^−1^ were attributed to changes in the position of the mirror, and a window of 0.025 s following the detected times was added to ensure full stabilization. The detected segments in the data during which the mirror moves were subtracted from further analysis (Supplementary Fig. 3).

### Data analysis

#### Conversion of the data into physical units

Data were digitized at a sampling rate fs=2500 Hz, and saved to a disk. All further processing of the data was done with Matlab (Mathworks). Using the calibration parameters previously obtained, the total extension of the tether, *x*, and the force acting on it, *F*, were calculated. From the extension *vs*. time traces, we identified the sections in the data containing SD polymerization. Then, the number of bp polymerized during SD and PE activity (*N*_*SD*_ and *N*_*PE*_, respectively), were calculated as:

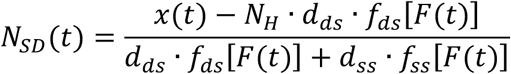

and

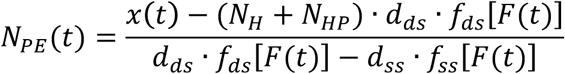

where *N*_*H*_=1200 is the number of bp in the dsDNA handles, *N*_*HP*_ is the number of bp in the hairpin, *d*_*ds*_ = 0.34 nm the rise per bp for dsDNA and *d*_*ss*_ = 0.66 nm the rise per base for ssDNA(39). *f*_*ds*_ and *f*_*ss*_ are functions describing the extension-over-contour ratio for dsDNA and ssDNA, respectively, as a function of the applied force. For the dsDNA parts, we used an extensible worm-like-chain (eWLC) model, and for the ssDNA parts, a WLC model(40). The persistence length was experimentally determined, for each molecule probed, by fitting force-extension curves.

#### Pause-free velocity calculation

The original, 2500 Hz data was low-pass filtered with a 3^rd^ order Butterworth filter with a cutoff *f*_*C*_ = 0.5 *Hz*, and the residency time in 1 bp windows, *τ*, was calculated. Data points corresponding to *τ* > *med*(*τ*) + 5 ∙ *madd*(*τ*), where *med*(*τ*) and *mad*(*τ*) are the median and the median absolute deviation of *τ*, respectively, were considered as belonging to pauses. Pauses shorter than 1s and polymerization bursts shorter than 2 bp, were discarded. The performance of the pause-detection scheme was tested using simulated traces(34), revealing ≤2% of false-negatives and ≤4% of false-positives across all the conditions tested(34). The pause free velocity was calculated as the mean instantaneous velocity from the segments where the enzyme in not paused.

#### Pause-density calculation

We previously showed that, under our experimental conditions, “pauses” longer than 20 s correspond to dissociation events followed by re-initiation(34). Hence, the pause density was calculated by counting the total number of pauses in the data set that had a duration shorter that 20 s, and then dividing by the total number of bp polymerized in the data set.

#### Fitting of the data

*V*_*max*_ and K_M_, as a function of force, and *PD*_1_, as a function of force and [dNTP], were globally fitted to their respective expressions, using Matlab’s *Global Search*, a scatter-search based global optimization method, running the solver *fmincon*. Confidence intervals for the fitted parameters were calculated by bootstrapping.

## RESULTS

### Processive polymerization by M-MLV Reverse Transcriptase is force- and [dNTP]-dependent

We followed SD polymerization by RT on a 265 bp DNA hairpin, that is attached to two ~ 600 bp dsDNA handles (Fig. 1A). Each handle harbors a tag (biotin and digoxigenin, respectively), allowing to tether the complete construct between two microspheres trapped in the two separate optical traps of a dual-trap high resolution optical tweezers (Fig. 1A). As shown previously(34), upon exposure of the complex to a mixture of RT an dNTPs, RT binds to the DNA construct by using the 3’-OH terminus of the dsDNA handle as a primer and engages in SD polymerization of the hairpin. SD polymerization results in an increase in the tether extension, by an amount equal to the addition of a single bp and a single nt for each catalytic cycle. With the known mechanical response of dsDNA and ssDNA, these extension changes can be translated into the number of bp polymerized (Figure 1B). We recently showed that polymerization is punctuated with numerous inactive phases, and that most of them represent backtracking of the polymerase(34). Here, we take advantage of our ability to reliably separate the pauses, and concentrate on the active phases of polymerization.

**Figure 1:**
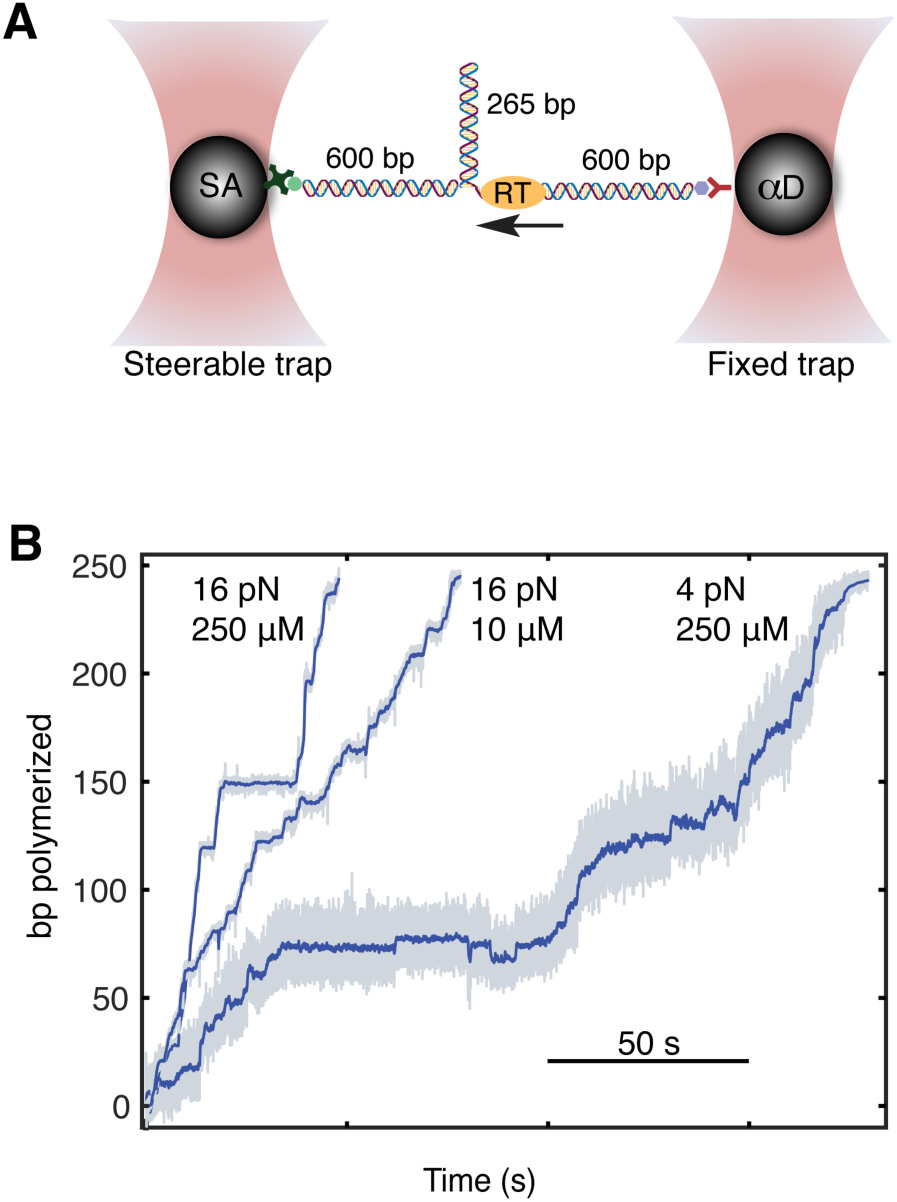
Following SD polymerization on a hairpin substrate under tension. (A) Experimental geometry. A 265 bp DNA hairpin is connected to two 600 bp dsDNA handles, and held under tension in a dual-trap optical tweezers. The 3’-OH terminus of one of the dsDNA handles serves as a primer for elongation (B) Time-traces of SD polymerization by RT. The extension of the tether is converted into the number of polymerized base pairs. Typical data is shown for different conditions of force and [dNTP].

Hence, we measure the pause-free velocity for different forces (4-16 pN) and a broad range of [dNTP] (0.5 – 250 μM). Fitting the data collected for each force indicates that, for all forces probed, the velocity follows Michaelis-Menten (MM) kinetics, *V* = *V*_*max*_([*dNTP*]/(*K*_*M*_ + [*dNTP*])) where *V* is the pause free velocity (Figure 2A). Moreover, the data shows that *V*_*max*_ is a sensitive function of the force (Fig. 2B), exhibiting a nearly 3-fold increase as the force is increased from 4 to 16 pN. The uncertainty in the determination of *K*_*M*_ is too large to determine its force-dependence, likely because of the difficulty of accurately characterizing the velocity at low force and [dNTP].

**Figure 2:**
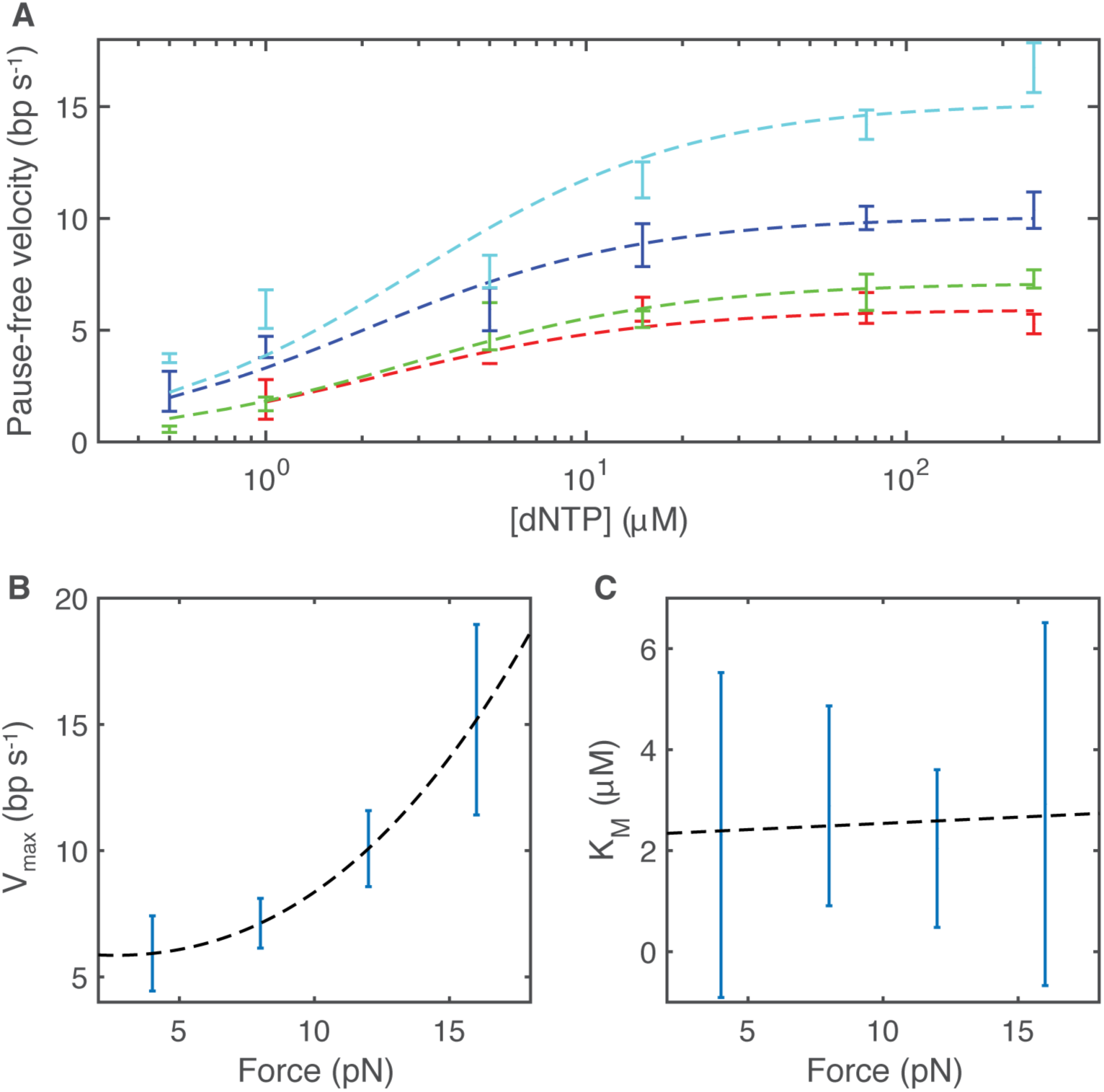
RT follows force-dependent Michaelis-Menten kinetics. (A) Pause free velocity for 4 (red), 8 (green), 12 (blue) and 16 (cyan) pN of force on the DNA. The pause free velocity was calculated, for a given force and [dNTP], by subtracting all pauses events in the polymerization trace. Data shown as mean ± sem. The number of traces measured is listed in Supplementary Table 6. The dashed lines show the results of fits to the Michaelis Menten equation. (B) *V*_*max*_ as a function of the applied force. Shown are the estimated values from the fit, and their 95% confidence interval. (C) *K*_*M*_ as a function of the applied force. Shown are the estimated values from the fit, and their 95% confidence interval.

### Mechano-chemical models for SD polymerization make distinct predictions for the dependence on force and [dNTP]

In order to extract from the data information about the translocation step during RTs polymerization reaction, we need to analyze how force affects the mechano-chemical cycle. Note, that previous studies have used an external force directly acting on a molecular motor to extract this information (see, for example, (13, 26, 41–43)). Here we show that, for an enzyme capable of performing SD polymerization, modulating the strength of the barrier imposed by the duplex ahead can fulfill a similar role.

Previous studies have delineated the elongation kinetic cycle of RT and other DNA polymerases as follows: Following binding of a new nucleotide, the complex undergoes a structural transition involving a hinged movement of the fingers domain and other smaller movements, from an “open” conformation which allows binding of nucleotides and, possibly, movement of the polymerase on the DNA, into a “closed” conformation that ensures proper positioning of the 3’ end of the primer, the template and the incoming nucleotide as if they were already in a dsDNA conformation. Next, a phosphodiester bond is formed, followed by a second structural transition (“opening” of the fingers). Finally, the pyrophosphate is released. Since with our experimental conditions PPi release is an irreversible transition ([PPi]≈ 0), this cycle can be simplified to include a nucleotide binding step, a catalysis step and a PPi release step, as follows(44, 45):

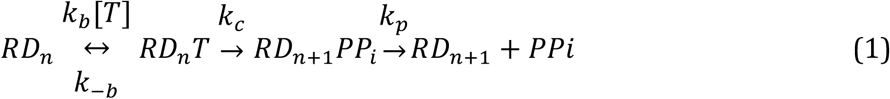

Here, *R* represents the RT enzyme, *D* is the DNA strand being synthesized, of length *n*, and *T* the incoming dNTP. However, being RT a processive enzyme that can perform successive incorporations without dissociation, there must be, in addition to the chemical steps described above, also a mechanical step of translocation, where RT moves by one bp on its template strand. As detailed above, this mechanical step may be located at different positions in the above cycle, and the mechanism underlying this step may correspond to either a PS or a BR model. Importantly, for both mechanisms, the end point of the reaction is progress along a chemical as well as a mechanical reaction coordinate. However, this is achieved via different pathways: For the PS mechanism translocation is tightly coupled to a specific chemical step, such us PPi release, and therefore can be represented by a diagonal movement in the two-dimensional energy landscape (Fig. 3C,D). For BRs, the movement between the pre- and post-translocation positions precedes the chemical step and the system moves in two orthogonal steps in its energy landscape (Fig. 3A,B): first along the mechanical coordinate, and then along the chemical one. This indicates that a PS mechanism should be integrated into the mechano-chemical cycle by considering a specific chemical step as a mechano-chemical step. For example, if translocation takes place with release of PPi, via a PS mechanism, we will simply consider *k*_*p*_ to be a step that includes both PPi release *and* translocation. Alternatively, formulating a BR model requires an additional, purely mechanical step to be added to the cycle, preceding the chemical step that rectifies the spontaneous fluctuations (the “pawl” of the ratchet). Hence, if translocation takes place via a BR rectified by PPi release, we add a pure translocation transition, with forward and backwards rates *k*_*t*_, *k*_*-t*_, between *k*_*c*_ and *k*_*p*_.

**Figure 3:**
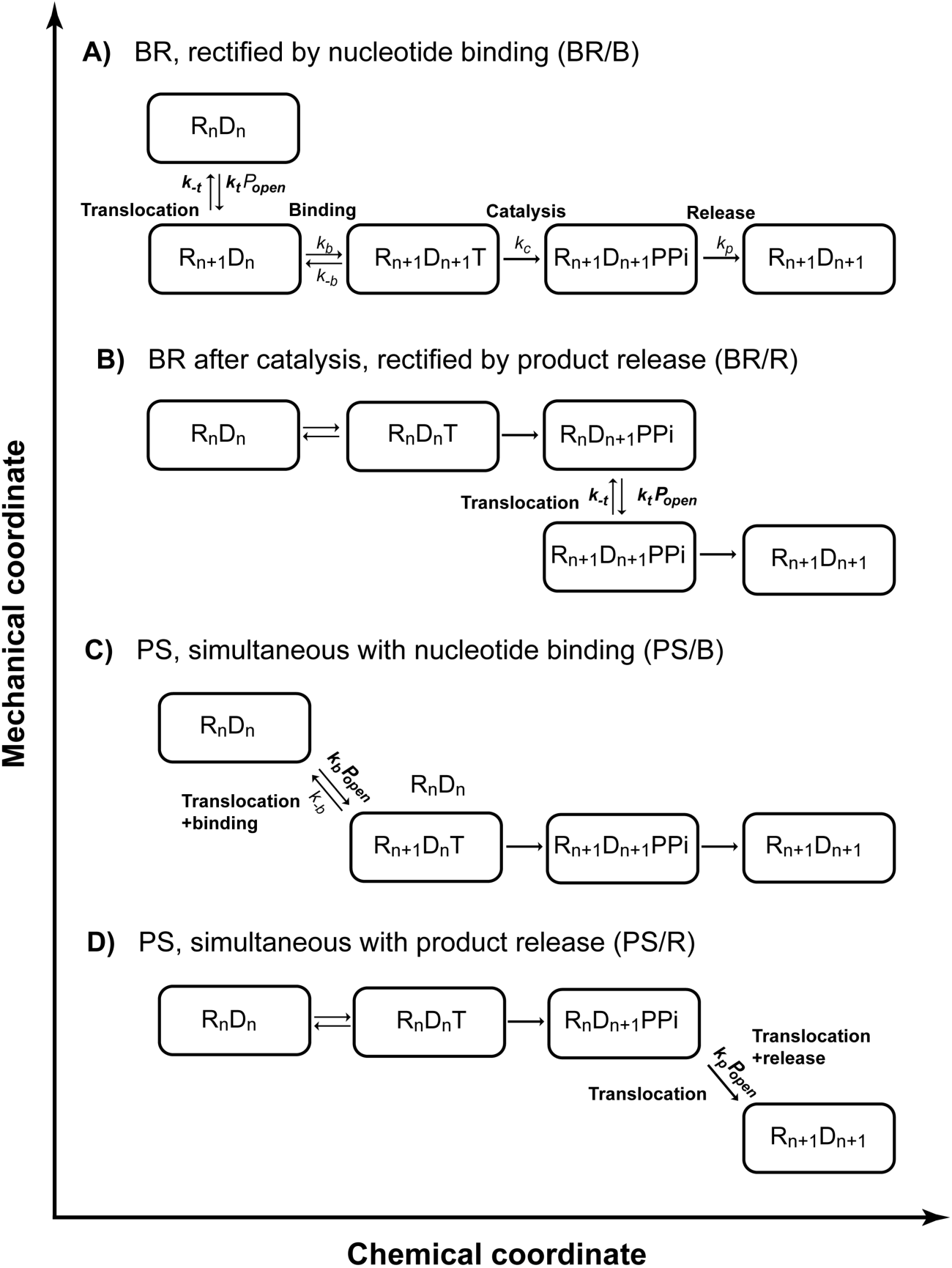
Different mechano-chemical schemes can incorporate a translocation step. Schematic description of the different mechano-chemical models used to predict the force- and [dNTP] dependence of the pause-free velocity. (A) A BR model, rectified by dNTP binding. (B) A BR model, where dNTP release locks the post-translocation state. (C) A PS model, where translocation is concomitant with dNTP binding. (D) A PS model, with translocation occurring together with release of dNTP.

In the special case of SD polymerization, when unwinding of the substrate is coupled to polymerization, translocation requires an open fork, i.e. a ssDNA region equal to or greater than the enzyme’s step size. Hence, we treat the open fork as a “substrate” of the translocation reaction and write the translocation step as 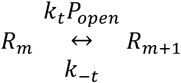, where *P*_*open*_ is the “concentration” of open forks, i.e. the probability for the fork to be open by at least the size of the enzyme’s step. As shown in the Supplementary Discussion, *P*_*open*_ depends on the free-energy difference between open and closed states, and is modulated by an external force applied on the hairpin, which stabilizes the open state. In addition, if the enzyme is not completely passive there will be some degree of destabilization of the closed state, given by a change in the free energy ∆*G*_*RT*_. Hence, both the force and ∆*G*_*RT*_ can modulate the translocation step. However, how this affects the overall rate of polymerization depends on the specific model for the mechano-chemical coupling that describes RT.

The first mechano-chemical model we consider is a BR where the rectification is provided by dNTP binding (BR/B model, Fig. 3A), as was previously suggested(28, 46), and demonstrated for other polymerases(26, 30). Such model can be described by the following reduced scheme:

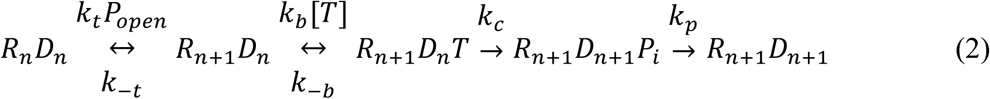

Previous studies have suggested that the rate limiting step in the cycle of DNAPs is the conformational change required for catalysis(2, 45, 47–49), and that both pyrophosphate release and translocation(2, 48, 50) are fast. This is also the case for RT, at least for its DNA-dependent synthesis activity(51–53). Hence, assuming that catalysis is the rate limiting step, i.e. k_c_ ≪ k_p_, k_-b_, k_-t_, we can show (Supplementary Information) that the polymerization velocity follows MM kinetics with 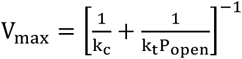 and 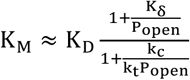, where we have introduced the notation 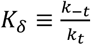 and 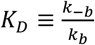. Three more models are considered (Fig. 3B-D and Supplementary Discussion): In a second BR model, the enzyme alternates between pre- and post-translocation after catalysis, and is rectified by the release of PPi (BR/R model). The last two additional models invoke a PS mechanism: In the first, translocation is coupled with binding of dNTP (PS/B model), and in the second it is coupled to the release of PPi (PS/R model). All the models predict MM kinetics, with parameters *V*_*max*_ and *K*_*M*_ that can be explicitly calculated (Supplementary Discussion, and Supplementary Tables 4). Notably, the models predict a different force dependence for the kinetic parameters, *V*_*max*_ and *K*_*M*_, as summarized in Table 1.

**Table 1:**
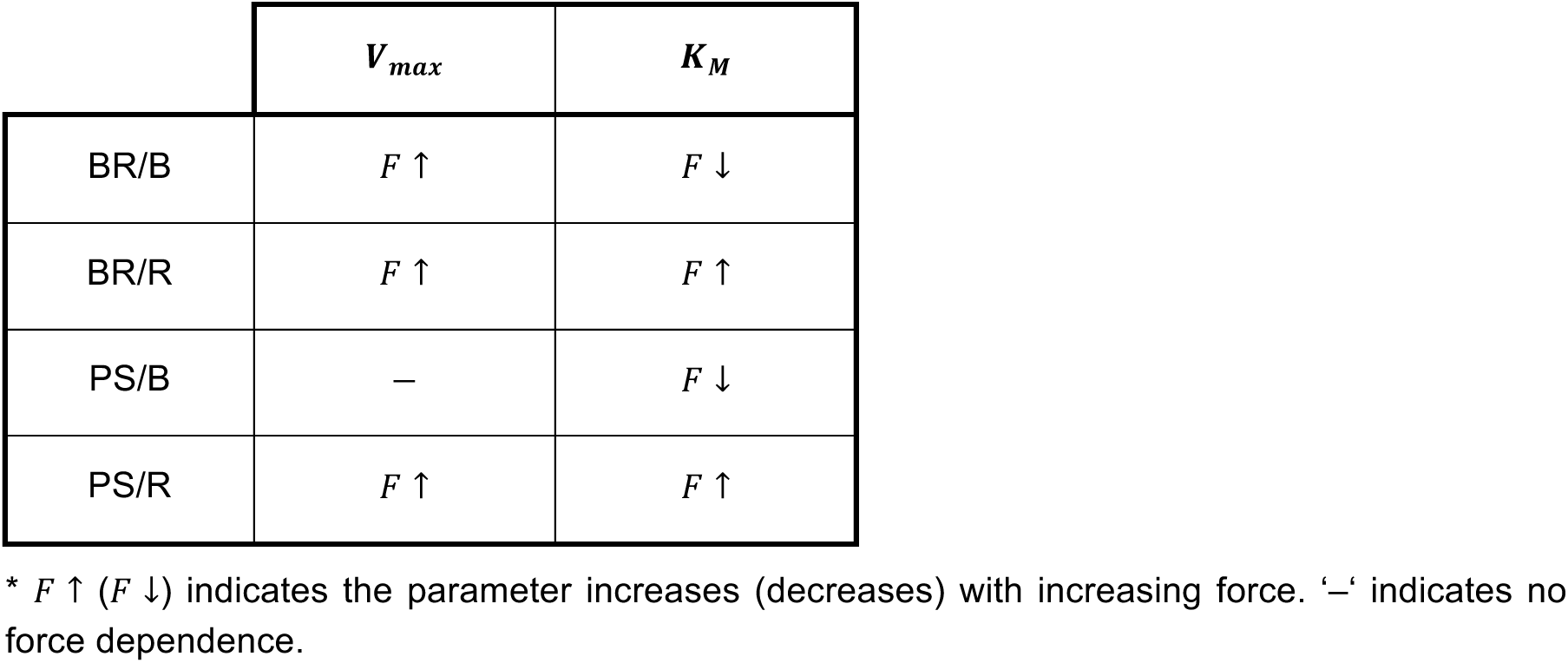
Force dependence of the Michaelis-Menten constants for the models tested.

As these models have different predictions for the dependence of the kinetic parameters on *p*_*open*_ and therefore on the applied force and ∆*G*_*RT*_, our measurements should help us elucidate which is the appropriate model to describe RT. In fact, they immediately rule out one of the models analyzed, PS/B, which predicts a force-independent *V*_*max*_, in disagreement with the monotonically-increasing dependence shown in Fig. 2B. The three other models we consider all predict a *V*_*max*_ that increases with force, and are consistent with the measurements. Unfortunately, the uncertainty in the determination of *K*_*M*_ is very large, preventing a significant discrimination between these models, which have different predictions for the force dependence of *K*_*M*_. However, as we show below, RT’s pausing phenotype will help us clarify this point.

### The mechano-chemical models predict the force- and [dNTP]-dependence of the pause density, and provide a constrain on the mechanism of polymerization

In our previous work, we showed that processive SD polymerization by M-MLV RT is punctuated by pauses, and that most of these pauses are backtracking events, at which the enzyme loses alignement with the 3’OH terminal of the primer. Notably, although pauses are off-pathway from the main polymerization cycle, a complete characterization of RT’s mechano-chemical cycle must incorporate the steps leading to the paused state. Hence, using the kinetic models of Fig. 3, we develop expressions for the Pause Density (PD), defined as the number of pauses per enzymatic cycle or, equivalently, the number of pauses per polymerized bp. We define *PD*_*j*_ as the probability, per cycle, to enter a pause from state *j*, where *j* is a number characterizing the position of a state in the cycle. So, for example, while state 1 is always the pre-translocated enzyme with no dNTP bound, the identity of state 2 can vary: for model BR/B is the post-translocation state, with no dNTP, and for model PS/B is the post-translocated enzyme with dNTP bound. *PD*_*j*_ can be calculated by examining the rate of entering the paused state from state *j* vs. the rate of productively continuing the cycle. The latter is given by 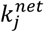 and therefore:

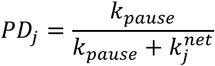

where *k*_*pause*_ is the intrinsic rate of entering the pause. Hence, to calculate *PD*_*j*_ we calculate 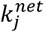 for a specific state of origin and a specific mechano-chemical model. For example, in the case of model BR/B, the pause can be entered from 4 different kinetic states (Fig. 3): i) the pre-translocation state, ii) the post-translocated state (before dNTP binding), iii) the dNTP bound, post-translocated state, and iv) the post-catalysis state, before PPi release. It can be shown (Supplementary Information) that under the assumptions that *k*_*c*_ is the rate limiting step,

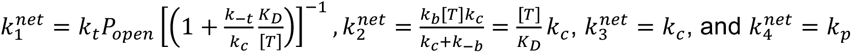

As can be seen, pauses originating from different states in the mechano-chemical cycle will exhibit a different force and [dNTP] dependence. Specifically, for the BR/B model, *PD*_1_ depends on both force and [dNTP], *PD*_2_ only on [dNTP], and *PD*_3,4_ are force- and [dNTP]-independent. In the same way, we calculate the predicted PDs for the rest of the models, and for each potential state of origin for the pauses within their mechano-chemical cycle (Supplementary Discussion, Supplementary Table 5). Their dependence on force and [dNTP] is summarized in Table 2.

**Table 2:** Force dependence of the pause-density for the models tested.

|  | Pause origin |  |  |  |
| --- | --- | --- | --- | --- |
|  | 1 | 2 | 3 | 4 |
| BR/B | $[T] \downarrow, F \downarrow$ | $[T] \downarrow$ | — | — |
| BR/R | $[T] \downarrow$ | — | $F \downarrow$ | — |
| PS/B | $[T] \downarrow, F \downarrow$ | — | — | |
| PS/R | $[T] \downarrow$ | — | $F \downarrow$ | |

\* $F \uparrow$ ( $F \downarrow$ ) indicates the parameter increases (decreases) with increasing force. $[T] \downarrow$ ( ) indicates the parameter increases (decreases) with [dNTP]. — indicates force- and [dNTP] independence

Our previous work showed that PD decreases as both force and [dNTP] are increased. Hence, the analysis presented above indicates that the only models compatible with the PD data are models BR/B and PS/B, and in both cases the pauses must originate from the first state in the cycle: the pre-translocated state. However, we showed above that PS/B is not compatible with the pause-free velocity data, since it predicts a force-independent Vmax. Hence, the only model which is compatible with both the velocity data and the pausing data, is model BR/B. We conclude that RT is a BR, rectified by dNTP binding, and that backtracking pauses are accessed from the pre-translocation state.

### RT is a relatively passive enzyme

Having established the architecture and mechanism of the mechano-chemical cycle of RT, we use our data to elucidate also the mechanism of strand displacement. Note, that since all the parameters affecting *P*_*open*_, with the exception of ∆G_*RT*_, are measured, we can consider *p*_*open*_, for our specific experimental setup, to be a function of ∆G_*RT*_ only (Supplementary Discussion). Hence, we fit the expressions we found for *V*_*max*_ and K_M_ (as a function of force) and *PD*_1_ (as a function of force and [dNTP]), to our experimental data (Supplementary Fig. 2), to find ∆G_*RT*_ and the microscopic rate constants governing the mechano-chemical cycle. As shown in Table 3, our data shows that during SD polymerization, RT is a moderately passive enzyme, destabilizing the fork only by ~ 1*k*_*B*_*T*.

**Table 3:** Microscopic rates obtained from fitting the pause-free velocity and PD

| $\Delta G_{RT}$ (k<sub>B</sub>T) | $k_c$ (s<sup>-1</sup>) | $k_t$ (s<sup>-1</sup>) | $k_{-t}$ (s<sup>-1</sup>) | $K_D$ (μM) | $k_{pause}$ (s<sup>-1</sup>) |
| --- | --- | --- | --- | --- | --- |
| $0.87 \pm 0.15$ | $58 \pm 37$ | $22 \pm 5$ | $220 \pm 145$ | $0.6 \pm 0.3$ | $0.4 \pm 0.1$ |

\* Shown are the estimated values and their 95% confidence interval

## DISCUSSION

In this work, we followed SD polymerization of M-MLV RT on a DNA template at the single molecule level. Separating the active phases of polymerization from the pauses induced by backtracking of the enzyme, we were able to characterize the pause-free velocity of RT in a broad range of mechanical and chemical conditions. We found that RT exhibits MM kinetics at all forces probed, and that the resulting *V*_*max*_ is a monotonically increasing function of the force applied on the duplex DNA. We postulated different kinetic models for the translocation, and derived the expected kinetic parameters for all of them. Comparing the predictions of these models with the force dependence of *V*_*max*_ enabled us to rule out one of the models. Next, we derived predictions for the pause density, for each of the kinetic models and for each of the possible states from which pauses are accessed, and compared the predictions of these models with the dependence of PD on force and [dNTP], as reported in our previous work(34). This allowed us to determine that only one model is compatible with both the pause-free velocity and the PD data, and to conclude that RT uses a BR mechanism, where binding of a new dNTP acts as the rectifier of translocation.

Interestingly, although our findings on the mechanism of translocation are in agreement with some previous studies, they disagree with others: Structural studies have suggested that HIV-1 RT translocation takes place by a PS model, and it was suggested that the conserved YMDD motif “buffers” the energy from dNTP binding as mechanical strain which powers translocation upon release(27). In contrast, a different study using site-specific footprinting and inhibition of HIV-1 RT by Foscarnet, supported a BR model rectified by dNTP binding(28, 29), similar to the one revealed in our work for M-MLV RT.

Being translocation an inherently mechanical process, its reliable characterization requires mechanical means, i.e. the application of force. Indeed, previous single molecule studies of RNA and DNA polymerases have used approaches similar to ours, based on globally fitting the data to a force-dependent model(13, 26). However, their approach was to study translocation under a force that directly opposes (or facilitates) the enzyme movement. This necessitates attaching the enzyme to one of the beads (or to a surface), a procedure that may affect its activity in unknown ways. Here we show that by considering the fork as an energetic barrier for translocation during SD synthesis, we can extract the same type of information as these previous works did for other enzymes. Therefore, we suggest that this approach may be of use to study the mechano-chemistry of other SD competent polymerases.

The retroviral genome is rich in secondary structure, with the strongest structural motifs present in HIV-1 being the TAR elements(14). Our results indicate that the destabilization of the fork by RT is very small, ~1 kBT, and are consistent with previous results for HIV-1 RT(32). This relative “passiveness” is what makes the velocity of RT highly sensitive to the presence and nature of a duplex. Recently, we showed that SD activity by RT is dominated by its pausing kinetics, where inactive states during processive polymerization are the result of backtracking of the enzyme, and are modulated by the strength of the duplex ~8 bp ahead of the fork. Here, we showed that these backtracking pauses are accessed from the pre-translocation state, in line with backtracking of RNAP(30). Together, the kinetic competition between elongation and backtracking, and the fact that both are modulated by the template’s secondary structure, suggest that backtracking can serve to amplify and diversify the regulatory effect of secondary structures on the polymerization, thus providing a mechanical basis for the regulation of RT by conserved structural motifs.

## SUPPLEMENTARY DATA

Supplementary Data are available at NAR online.

## Author Contributions

A.K. designed and supervised the research. O.M. and H.K. performed the experiments. S.R. prepared reagents. O.M. and A.K. designed and built the optical tweezers setup. O.M., H.K. and A.K. analyzed the data. A.K. wrote the paper.

